# Common variation at the LRRK2 locus is associated with survival in the primary tauopathy progressive supranuclear palsy

**DOI:** 10.1101/2020.02.04.932335

**Authors:** Edwin Jabbari, Manuela M.X. Tan, Regina H. Reynolds, Kin Y. Mok, Raffaele Ferrari, David P. Murphy, Rebecca R. Valentino, Owen A. Ross, Dennis W. Dickson, Safa Al-Sarraj, Steve M. Gentleman, Kieren S.J. Allinson, Zane Jaunmuktane, Janice L. Holton, Tamas Revesz, Thomas T. Warner, Andrew J. Lees, Mark R. Cookson, J. Raphael Gibbs, Jinhui Ding, Ruth Chia, Bryan J. Traynor, Sonja W. Scholz, Alexander Pantelyat, Coralie Viollet, Clifton L. Dalgard, Olga Pletnikova, Juan C. Troncoso, Adam L. Boxer, Gesine Respondek, Günter U. Höglinger, David J. Burn, Nicola Pavese, Alexander Gerhard, Christopher Kobylecki, P. Nigel Leigh, Alistair Church, Michele T.M. Hu, James B. Rowe, Mina Ryten, John Hardy, Maryam Shoai, Huw R. Morris

## Abstract

The genetic basis of variation in the rate of disease progression of primary tauopathies has not been determined. In two independent progressive supranuclear palsy cohorts, we show that common variation at the *LRRK2* locus determines survival from motor symptom onset to death, possibly through regulation of gene expression. This links together genetic risk in alpha-synuclein and tau disorders, and suggests that modulation of proteostasis and neuro-inflammation by LRRK2 inhibitors may have a therapeutic role across disorders.

## Introduction

Progressive supranuclear palsy (PSP) is a rapidly progressive neurodegenerative tauopathy. In the classical form, PSP-Richardson syndrome (PSP-RS), patients develop imbalance and frequent falls, bulbar failure and dementia, and survive a mean of 6.9 years from symptom onset [1]. More recently defined PSP subtypes, such as PSP-Parkinsonism and PSP-Progressive Gait Freezing, are associated with a slower rate of progression [2]. We have shown that the PSP phenotype is determined by variation at the *TRIM11/17* locus [3]. The pathology of PSP involves the deposition of insoluble hyperphosphorylated tau predominantly in neurons and astrocytes with involvement of sub-cortical and cortical brain regions. Recent work in animal models has shown that PSP-tau pathology is transmissible, in that human inoculate in mice leads to PSP-type pathology in the recipient which recapitulates the morphology and immunohistochemical features of the human tauopathy [4].

Genome-wide studies in neurodegeneration have focussed on case-control status which have provided powerful insights into the aetiology of neurodegenerative disease. In PSP, common variants at the *MAPT, MOBP, EIF2AK3* and *STX6* loci are associated with PSP risk [5]. However, therapeutic efforts focus on developing therapies which slow or halt disease progression following clinical diagnosis. The genetic determinants of clinical disease progression for common neurodegenerative diseases are largely unexplored and are likely to provide important clues to biology and effective therapy. Clinical disease progression relates to the sequential involvement of brain areas and systems. This may relate to differential neuronal susceptibility/resistance (cell-autonomous) or cell to cell spread (non-cell-autonomous) factors.

## Results

Here, we have used a Cox-proportional hazards survival model to identify genetic determinants of survival (from motor symptom onset to death) in 1,001 PSP cases of European ancestry derived from two independent cohorts (Data Supplement Table 1). Following genotype data quality control and imputation against the Haplotype Reference Consortium v1.1 panel, we found no association between progression and previously identified PSP risk variants (p>0.05) after adjusting for sex, age at motor symptom onset and the first three principal components. There was an association between the *TRIM11/17* locus and progression (rs564309, hazard ratio = 0.85, p=0.01), but this was conditional on PSP phenotype. We then conducted a survival genome-wide association study (GWAS) using PSP phenotype as an additional binary (PSP-RS or non-PSP-RS) covariate and analysed 4,817,946 common (minor allele frequency ≥1%) SNPs. In the whole-cohort analysis we found an association between variation at rs2242367 and PSP survival (GRCh37 chr12:40413698, p=7.5×10^-10^), HR 2.24) (Figure 1a). There appeared to be an additive allele / survival relationship (mean [standard deviation] disease duration in deceased cases: reference GG - 7.7 [3.10] years, AG - 6.9 [2.78] years, AA - 6.6 [2.74] years). The association between rs2242367 and survival was replicated in each cohort (Table 1). rs2242367 is not associated with PSP risk or PSP phenotype (Data Supplement Table 2). A conditional analysis that controlled for rs2242367 genotype did not show other independent association signals.

**Table 1:**
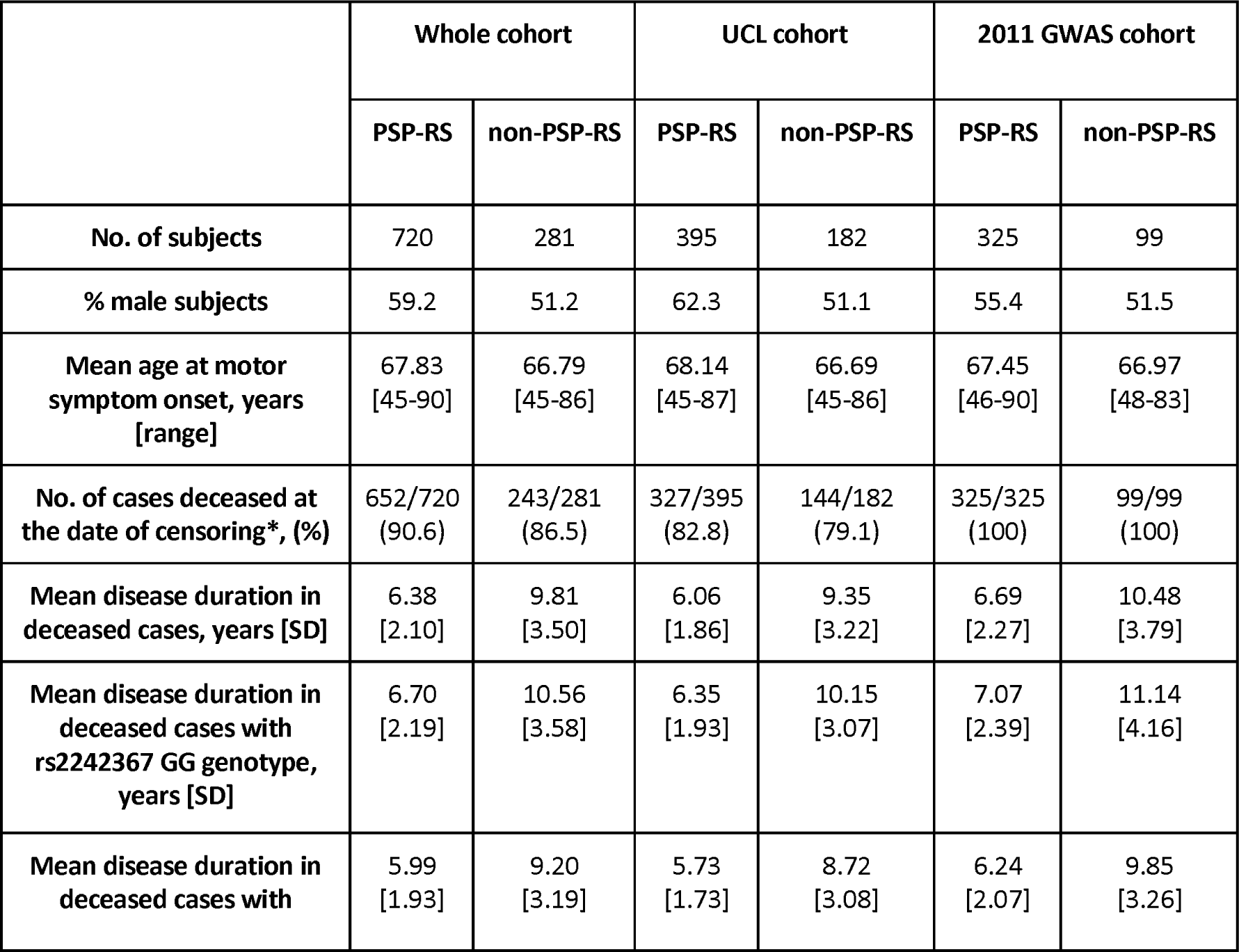
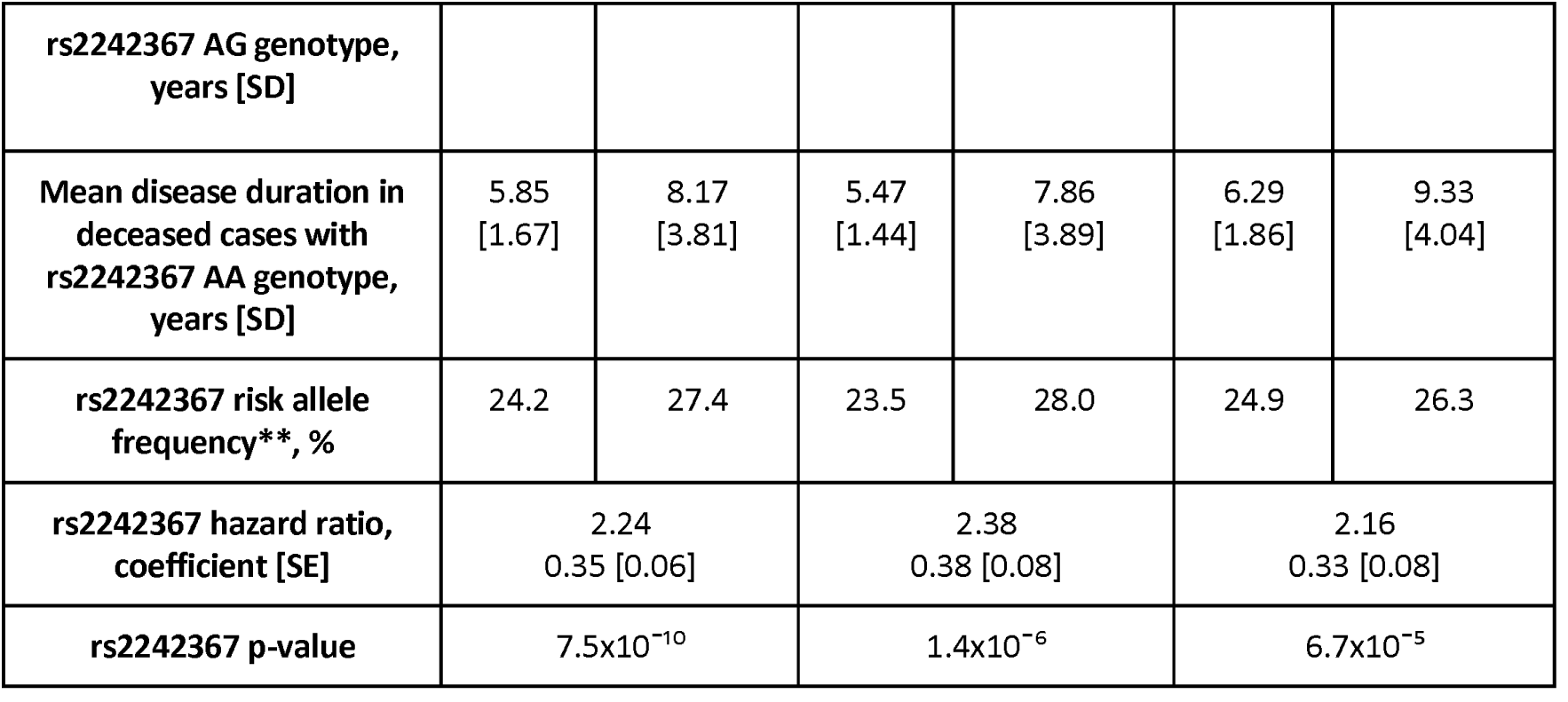
Clinical characteristics and rs2242367 association statistics from PSP survival GWAS, stratified by cohort and PSP phenotype

**Figure 1:**
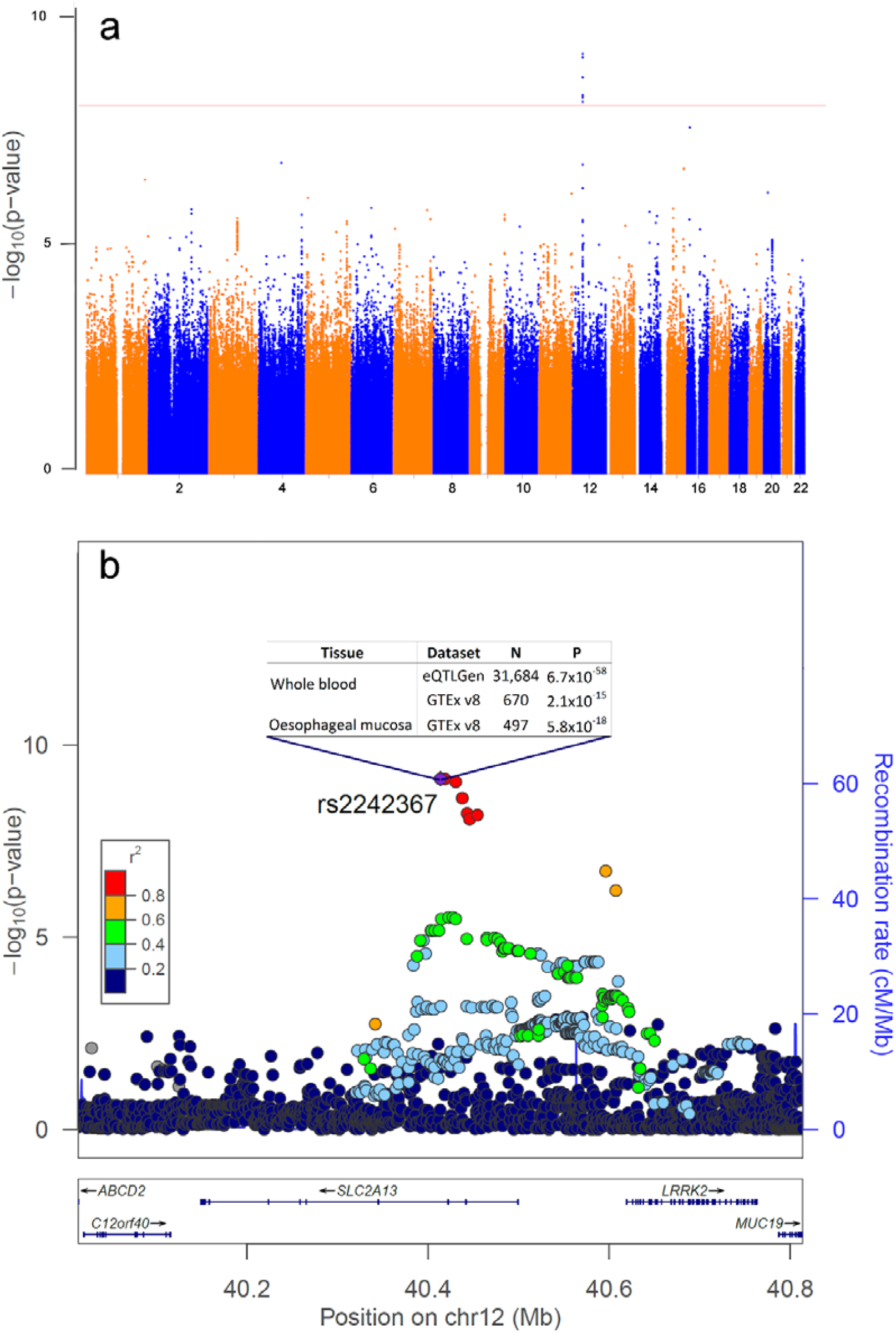
Manhattan and regional association plots, with supporting eQTL data, highlighting LRRK2 association with PSP survival. **a**, Manhattan plot of whole-cohort PSP survival GWAS, highlighting a genome-wide significant signal at chromosome 12. The red line indicates the Bonferroni-corrected threshold for genome-wide significance (p<1.0×10^-8^). **b**, Regional association plot of whole-cohort PSP survival GWAS, identifying rs2242367 as the lead SNP with associated eQTL signals for LRRK2. SNP positions, recombination rates and gene boundaries are based on GRCh37/hg19.

Clinical characteristics of cases included in the PSP survival GWAS. Results are accompanied by either standard deviation (SD) or standard error (SE) measurements. * = Date of censoring was 01/12/2019. ** = rs2242367 risk allele frequency in 7,692 reference controls of non-Finnish European ancestry was 28.2%, taken from https://gnomad.broadinstitute.org on 10/01/2020.

### Structure of the region

rs2242367 lies within intron 3 of SLC2A13, 190Kb from the common LRRK2 risk SNP for Parkinson’s disease (PD), rs76904798 (GRCh37 chr12:40614434, p=1.5×10^-28^, OR 1.15) [6]. The two variants are in r^2^ linkage equilibrium in non-Finnish European populations, but with a high D’ (r^2^=0.05, D’=0.91) (LDlink v3.8). The allele frequency of rs2242367 is 28% and rs75904798 is 13% (GnomAD v2.1.1) so the LRRK2 PD risk SNP likely defines a sub-haplotype of the ancestral PSP survival risk haplotype. This region also contains association signals for two immune/inflammatory disease associations, Crohn’s disease and Leprosy [7].

### Functional annotation

There are no coding variants in linkage disequilibrium with the lead SNP rs2242367 as defined by the region encompassed by variants with an r^2^>0.3 (Data Supplement Figure 1). We also explored the expression quantitative trait loci (eQTL) status of rs2242367 using multiple eQTL datasets via FUMA (https://fuma.ctglab.nl) (Methods). We found that rs2242367, and its associated tagging intronic SNPs, were strong eQTLs for increasing LRRK2 expression in whole blood and oesophagus in eQTLGen and GTEXv8 and datasets (Figure 1b, eQTLGen LRRK2 whole blood p=6.7×10^−58^). rs2242367 was also an eQTL for long intergenic non-coding RNA (lincRNA) RP11-476D10.1 (ENSG00000260943, p=3.3×10^−310^) and AC079630.4 (ENSG00000223914, p=5. ×10^−22^) in the eQTLGen whole blood dataset [8]. Additionally, we interrogated the North American Brain Expression Consortium (NABEC) dataset (213 human frontal cortex samples) and found no significant eQTL signals.

### WGS data

A subset of our cases (n=140) underwent whole-genome sequencing (WGS). In these cases, the allele frequencies of our significant SNPs had 100% concordance between WGS and chip-based imputed datasets. We ran the Cox-proportional hazards survival model on variants with a minor allele frequency ≥0.1% in our region of interest (GRCh38 chr12:39225001-41369277) and found 214 variants with a stronger association signal than rs2242367 (p<0.01). All of these were non-coding variants in linkage disequilibrium with rs2242367 (r^2^=0.30-0.35, D’=0.96-1.0) and had the same eQTL profile as rs2242367 (eQTLGen LRRK2 whole blood p<1.0×10^−20^).

## Discussion

We have shown that common variation at the *LRRK2* locus is a genetic determinant of PSP survival. *LRRK2* is an established major risk factor for PD with common and rare (G2019S) variants associated with disease [6]. G2019S can cause familial PD, although often with reduced penetrance. Recently, the *LRRK2* PD risk-associated SNP, rs76904798, has also been associated with the rate of progression of PD [9]. LRRK2 is expressed in multiple human tissues including brain and whole blood. In brain, it is expressed ubiquitously across all regions and is found in neurons, astrocytes, microglia and oligodendroglia [10]. Pathogenic mutations in *LRRK2* lead to phosphorylation of a subset of Rab proteins which have important roles in the formation and trafficking of intracellular vesicles [11]. This in turn may affect both proteostasis and the inflammatory response, which may be important in mediating disease progression in PSP.

There is also an established link between LRRK2 and tau pathology, which was reported in chromosome 12 linked PD families before the identification of the *LRRK2* gene [12]. In rare cases, *LRRK2* mutations have been identified in patients with PSP-tau pathology at post-mortem [13]. Recently, the link between LRRK2 and tau has been explored in cell and animal models, identifying dysregulation of actin and mitochondrial dynamics, and the impairment of tau degradation via the proteasome [14-16].

Our lead SNP, rs2242367, is an eQTL and a plausible explanation is that this effect is mediated through the control of LRRK2 expression. Data from our subset of cases with WGS validates the genotyped data used for our GWAS and suggests that the association signal at rs2242367 may be part of a larger haplotype block, paving the way for further studies using larger WGS datasets to clarify this.

The strongest evidence for the effect on expression comes from blood rather than bulk RNA analysis from brain. This may relate to the cell types present and sample size differences in the blood and brain expression datasets. In the PsychENCODE bulk RNA dataset, most of the power for detecting eQTL signals in the brain comes from astrocytes and neurons, with microglia representing only 5% of normalised cell fractions [17]. Alternatively, the predominant effect on expression in blood may relate to peripheral immune response-driven neuroinflammation [18] or be due to a specific effect in monocyte/microglial lineage cells. The advent of expression analysis in defined brain cell subpopulations will clarify this issue. Increased LRRK2 expression may result in a reactive microglia-induced pro-inflammatory state which drives ongoing accumulation of misfolded tau protein and clinical disease progression [19, 20]. This hypothesis is supported by *in vivo* positron emission tomography evidence of a pattern of microglial activation involving cortical regions and the basal ganglia that corresponds well with the known distribution of neuropathological changes [21, 22]. In mouse models, microglial inflammatory responses are attenuated by LRRK2 inhibition and this strategy is currently under investigation in PD as a disease-modifying therapy.

Our study paves the way for further functional studies assessing the impact of LRRK2 on tau aggregation and exploration of LRRK2 inhibition as a therapeutic approach in patients with tauopathies.

## Acknowledgements

This study was funded by grants from: the Medical Research Council to E.J. (548211); Parkinson’s UK to M.M.X.T. and M.T.M.H.; the Leonard Wolfson Doctoral Training Fellowship in Neurodegeneration to R.H.R.; NINDS Tau Center without Walls Program (U54-NS100693) to R.R.V., O.A.R. and D.W.D.; Department of Health’s NIHR Biomedical Research Centre’s funding scheme to Z.J.; the MSA Trust, the MSA Coalition and Fund Sophia, managed by the King Baudouin Foundation to J.L.H.; CBD Solutions and NIHR Queen Square Biomedical Research Unit in Dementia based at University College London Hospitals, University College London to T.R.; the Deutsche Forschungsgemeinschaft (DFG, EXC 2145 SyNergy – ID 390857198, HO2402/18-1 MSAomics), the German Federal Ministry of Education and Research (BMBF, 01KU1403A EpiPD; 01EK1605A HitTau), the NOMIS foundation (FTLD project), the German Center for Neurodegenerative Diseases (DZNE) to G.U.H.; the Wellcome Trust (103838), the Cambridge Centre for Parkinson-plus and NIHR Cambridge Biomedical Research Centre to J.B.R.; the Medical Research Council (MR/N026004/1), Wellcome Trust (202903/Z/16/Z), Dolby Family Fund and NIHR Biomedical Research Centre at University College London Hospitals NHS Foundation Trust and University College London to J.H.; anonymous donor to M.S.; the PSP Association to H.R.M. The PROSPECT study is supported by grants from the PSP Association, CBD Solutions, the MSA Trust and NIHR UCLH Biomedical Research Centre. Queen Square Brain Bank is supported by the Reta Lila Weston Institute for Neurological Studies and the Medical Research Council. Cambridge Brain Bank is supported by the NIHR Cambridge Biomedical Research Centre. The brain bank at Mayo Clinic in Jacksonville is supported by CurePSP and the Tau Consortium. This research was supported in part by the Intramural Research Program of the National Institutes of Health (National Institute on Aging, National Institute of Neurological Disorders and Stroke; project numbers: ZIA-AG000935, ZIA-NS003154). Tissue samples and clinicopathological information were provided by the Johns Hopkins Morris K. Udall Center of Excellence for Parkinson’s disease Research (NIH P50 N38377) and the Johns Hopkins Alzheimer Disease Research Center (NIH P50 AG05146). This study used the high-performance computational capabilities of the Biowulf Linux Cluster at the National Institutes of Health, Bethesda, Maryland, USA (http://biowulf.nih.gov). This work was supported by the UK Dementia Research Institute which receives its funding from DRI Ltd, funded by the UK Medical Research Council, Alzheimer’s Society and Alzheimer’s Research UK.

## Author contributions

E.J. and H.R.M. conceived and headed the project. E.J. carried out the statistical analysis of genetic data. M.S. supervised the statistical analysis of genetic data and created Figure 1. D.P.M. assisted with the quality control of genetic data. E.J., M.M.X.T., K.Y.M. and R.F. contributed to the genotyping of DNA samples. R.H.R., M.R.C., J.R.G. and M.R. contributed to the interpretation of eQTL results. J.D., R.C., B.J.T., S.W.S., V.C. and C.L.D. generated WGS data at NIH. E.J. was responsible for the phenotypic characterisation of cases and wrote the first draft of the manuscript. Critical revision of the manuscript was provided by E.J., A.J.L., B.J.T., A.L.B., J.B.R., M.R., J.H., M.S. and H.R.M. E.J., R.R.V., O.A.R., D.W.D., S.A.S., S.M.G., K.S.J.A., Z.J., J.L.H., T.R., T.T.W., S.W.S., A.P., O.P., J.C.T., G.R., G.U.H., D.J.B., N.P., A.G., C.K., P.N.L., A.C., M.T.M.H., J.B.R. and H.R.M. played a role in the acquisition of blood/tissue samples for DNA extraction and/or the pathologic and phenotypic characterisation of cases. All authors approved the final version of the manuscript.

## Competing interests

Professor Huw Morris is employed by UCL. In the last 24 months he reports paid consultancy from Biogen, UCB, Abbvie, Denali, Biohaven, Lundbeck; lecture fees/honoraria from Biogen, UCB, C4X Discovery, GE-Healthcare, Wellcome Trust, Movement Disorders Society; Research Grants from Parkinson’s UK, Cure Parkinson’s Trust, PSP Association, CBD Solutions, Drake Foundation, Medical Research Council. Professor Morris is a co-applicant on a patent application related to C9ORF72 - Method for diagnosing a neurodegenerative disease (PCT/GB2012/052140).

## Methods

### Study samples

Cases with a neuropathological diagnosis of PSP were identified from UK, US and German brain banks. Each brain bank had approved ethics to cover the analysis of tissue/DNA samples, and associated clinical data, for research projects. Cases with a clinical diagnosis of a PSP were identified from the Progressive Supranuclear Palsy-Cortico-Basal Syndrome-Multiple System Atrophy (PROSPECT) study, a UK-wide longitudinal study of patients with atypical parkinsonian syndromes (Queen Square Research Ethics Committee 14/LO/1575). A subset of deceased PROSPECT study PSP cases had post-mortem neuropathological confirmation of diagnosis at Queen Square (by Z.J., J.L.H. and T.R.) and Cambridge (by K.S.J.A.) brain banks.

### Clinical characterisation

All PSP cases were assigned a Movement Disorder Society PSP diagnostic criteria phenotype based on clinical features that were present in the first three years from motor symptom onset, using consensus criteria on application of the diagnostic criteria [23]. Cases were subsequently stratified into PSP-RS and non-PSP-RS groups, where non-PSP-RS consisted of PSP-parkinsonism and PSP-progressive gait freezing phenotypes. This approach was applied retrospectively using detailed case notes for brain bank cases (UCL cohort cases by E.J.; 2011 GWAS cohort cases by D.W.D., G.R. and G.U.H). PROSPECT study PSP cases were assigned a baseline PSP phenotype on entry to the study by E.J. using the same criteria as above. Of note, all included PROSPECT study PSP cases were at least three years into their disease course (from motor symptom onset) and fulfilled at least “possible” criteria for a PSP phenotype. In addition, the following clinical data was collected for all cases: sex; age at motor symptom onset; disease duration from motor symptom onset to death, or motor symptom onset to date of censoring (01/12/2019) for living PROSPECT study PSP cases.

### Genotype data

All brain bank cases had DNA extracted from frozen brain tissue (cerebellum or frontal cortex). PROSPECT study cases had DNA extracted from blood. Subsequently, DNA samples from all cases underwent genotyping using the Illumina NeuroChip (UCL cohort cases) and the Illumina Human 660W-Quad Infinium chip (2011 GWAS cohort cases) [5, 24]. Standard genotype data quality control steps were carried out, including sex checking, identity by descent and a principal components analysis (PCA) to exclude all cases of non-European ancestry [25]. All cases were screened for known *MAPT, LRRK2 and DCTN1* mutations covered by both genotyping platforms and excluded if found to be positive. The two datasets were merged and approximately 230K SNPs which were present on both genotyping platforms were extracted and used for SNP imputation via the Sanger Imputation Service. The following post-imputation data quality control steps were used to filter out SNPs: INFO score <0.70, posterior probability <0.90, missingness (geno) >0.05, Hardy-Weinberg equilibrium p-value >1.0×10^−7^ and minor allele frequency <0.01. Imputed SNP positions were based on Genome Reference Consortium Human 37/human genome version 19 (GRCh37/hg19).

### Statistical analyses

In total, 32 cases were excluded from this study due to either insufficient clinical data or having at least one of the above genotype data exclusion criteria, including one case that was found to have a previously described *MAPT* L284R mutation, resulting in 1,001 cases of European ancestry for analyses (Supplementary Table 1). Following post-imputation data quality control, 4,817,946 common (minor allele frequency ≥1%) SNPs for used for analyses. We used a cox-proportional hazards survival GWAS model that adjusted for sex, age at motor symptom onset, PSP phenotype (PSP-RS or non-PSP-RS) and the first three principal components derived from PCA. All reported significant SNPs had a Cox.zphP value >0.05, indicating that the model adheres to the assumption of proportional hazards. These analyses and the creation of Figure 1a was carried out in R version 3.3.2. Figure 1b was created using LocusZoom version 0.10. The Bonferroni-corrected threshold for genome-wide significance was set at p<1.0×10^−8^. We used GCTA-COJO to assess the independence of GWAS signals [26].

### eQTL data

We assessed the eQTL signals of all significant SNPs from our GWAS using FUMA, a platform to annotate, prioritise, visualise and interpret GWAS results [27]. The following eQTL data sources are used as part of FUMA’s eQTL analysis, with a more detailed description of tissue types and sample sizes available at https://fuma.ctglab.nl/tutorial#snp2gene: GTEx version 6, Blood eQTL browser, BIOS QTL browser, BRAINEAC, GTEx version 7, MuTHER, xQTLServer, CommonMind Consortium, eQTLGen, PsychENCODE, DICE, van der Wijst et al. scRNA eQTLs and GTEx version 8. Additionally, the NABEC eQTL dataset was analysed (dbGaP Study Accession: phs001300.v1.p1).

### WGS data

A subset (n=140) of PROSPECT study PSP cases of European ancestry underwent WGS at the National Institutes of Health (NIH). Blood-derived DNA samples were diluted to a target concentration of 50 ng/μl in Low-EDTA TE buffer (Quality Biological). PCR-free, paired-end, non-indexed libraries were constructed by automated liquid handlers using the Illumina TruSeq chemistry. DNA libraries were denatured and diluted to a final DNA concentration of 20 pM, followed by ‘single library’ - ‘single lane’ clustering on patterned flow cells using the Illumina cBot system. The libraries were sequenced on an Illumina HiSeq X Ten sequencer (v2.5 chemistry, Illumina) using 150 bp, paired-end cycles.

Raw genome data in FASTQ file format were transferred to Google Cloud Storage. Paired-end sequences were processed in accordance with the pipeline standard developed by the Centers for Common Disease Genomics (CCDG; https://www.genome.gov/27563570/). Sequence alignment to the human reference genome (GRCh38) was achieved using the Burrows-Wheeler Aligner (BWA-MEM) [28] and was followed by post-alignment processing for variant discovery using the Genome Analysis Toolkit Best Practices pipeline (GATK v4.1.2.0) [29]. The average sequencing read-depth after filtering by alignment quality was 35x. Single nucleotide and short indel variants were called from the processed WGS data using GATK HaplotypeCaller to generate individual gVCF files for each subject and followed by joint genotype calling of merged gVCFs to generate a multi-sample VCF file. Standard variant filtering was then performed using GATK4 Variant Quality Score Recalibration (VQSR) tools VariantRecalibrator and ApplyRecalibration.

For sample-level quality control checks, we first excluded genomes with a high contamination rate (> 5% based on VerifyBamID freemix metric) [30]. Next, we converted the multi-sample VCF-file to a PLINK 2.0 binary file (cog-genomics.org/plink/2.0) [31] and removed samples with an excessive heterozygosity rate (exceeding +/- 0.15 F-statistic), a sample call rate ≤ 95%, discordance between reported sex and genotypic sex, and duplicate samples (determined by pi-hat statistic). For variant-level quality control, we excluded variants with an overall missingness rate of > 5%, non-autosomal variants and variants that significantly departed from Hardy-Weinberg equilibrium (p<1.0×10^-10^).

### Data availability

The data that support the findings of this study are available in anonymised format by request of a qualified investigator to the corresponding authors.

## Supplementary Data

**Supplementary Table 1:** PSP sample sources

| Sample source | Number of samples | % of samples with post-mortem confirmation of PSP pathology |
| --- | --- | --- |
| **UCL PSP cohort** |  |  |
| Queen Square Brain Bank, UK | 248 | 100 |
| PROSPECT study, UK | 180 | 11* |
| Cambridge Brain Bank, UK | 54 | 100 |
| Johns Hopkins Brain Bank, USA | 37 | 100 |
| MRC London Neurodegenerative Diseases Brain Bank, UK | 25 | 100 |
| Multiple Sclerosis and Parkinson's Brain Bank, UK | 21 | 100 |
| Newcastle Brain Bank, UK | 12 | 100 |
| **2011 GWAS cohort** |  |  |
| Mayo Clinic Brain Bank, USA | 340 | 100 |
| Munich Brain Bank, Germany | 84 | 100 |

\*=74/180 PROSPECT study cases were deceased at the date of censoring. 20/74 had a post-mortem examination, of which 20/20 (100%) had a pathological diagnosis of PSP.

**Supplementary Table 2:** rs2242367 association statistics in PSP case-control and phenotype GWAS’

|  | PSP case-control GWAS* | PSP phenotype GWAS** |
| --- | --- | --- |
| **Odds ratio** | 0.95 | 0.80 |
| P-value | 0.35 | 0.21 |

\*=Summary statistics from phase 1 (pathologically confirmed PSP cases) Hoglinger et al. [5]
\*\*=Summary statistics from Jabbari et al. [3]

**Supplementary Figure 1:**
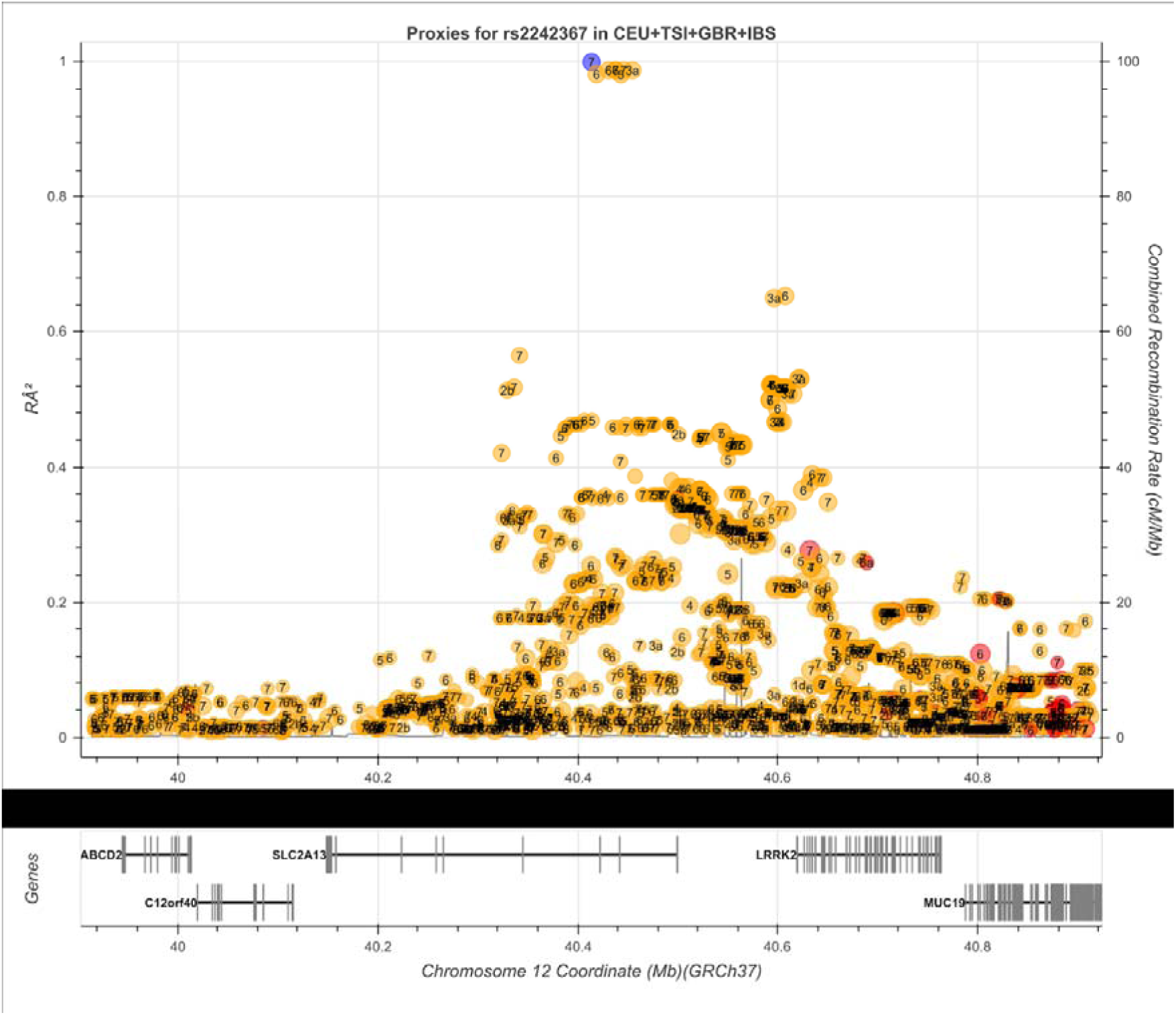
LD structure around rs2242367. LD plot from LDlink (https://ldlink.nci.nih.gov/) highlighting that there are no coding variants in linkage disequilibrium with the lead SNP rs2242367, as defined by the region encompassed by variants with an r^2^>0.3. Blue dot = lead SNP rs2242367, yellow dots = non-coding SNPs, red dots = coding SNPs. Regulatory potential of each SNP (based on RegulomeDB scores - http://www.regulomedb.org/help#score) indicated by numerical value between 1 (high) to 7 (low).

## References

1. Coyle-Gilchrist, I.T.S. et al. Neurology 86, 1736–1743 (2016).

2. Jabbari, E. et al. JAMA Neurol. 77, X–X (2020).

3. Jabbari, E. et al. Ann. Neurol. 84, 485–496 (2018).

4. Clavaguera, F. et al. Proc. Natl. Acad. Sci. 110, 9535–9540 (2013).

5. Höglinger, G.U. et al. Nat. Genet. 43, 699–705 (2011).

6. Nalls, M.A. et al. Lancet Neurol. 18, 1091–1102 (2019).

7. de Lange, K.M. et al. Nat. Genet. 49, 256–261 (2017).

8. Võsa, U. et al. bioRxiv 447367, doi: https://doi.org/10.1101/447367 (2018).

9. Iwaki, H. Neurol. Genet. 5, e348 (2019).

10. Miklossy, J. et al. J. Neuropathol. Exp. Neurol. 65, 953–963 (2006).

11. Alessi, D.R. & Sammler, E. Science 360, 36–37 (2018).

12. Funayama, M. et al. Ann. Neurol. 51, 296–301 (2002).

13. Sanchez-Contreras, M. et al. Mov. Disord. 32, 115–123 (2017).

14. Bardai, F.H. et al. PLoS Biol. 16, e2006265 (2018).

15. Guerreiro, P.S. et al. Mol. Neurobiol. 53, 3124–3135 (2016).

16. Nguyen, A.P.T. et al. Hum. Mol. Genet. 27, 120–134 (2018).

17. Wang, D. et al. Science 362, eaat8464 (2018).

18. Cao, W. & Zheng, H. Mol. Degen. 13, 51 (2018).

19. Maphis, N. et al. Brain 138, 1738–1755 (2015).

20. Moehle, M.S. et al. J. Neurosci. 32, 1602–1611 (2012).

21. Gerhard, A. et al. Mov. Disord. 21, 89–93 (2006).

22. Passamonti, L. et al. Neurology 90, e1989–1996 (2018).

## Methods references

23. Grimm, M.J. et al. Mov. Disord. 34, 1228–1232 (2019).

24. Blauwendraat, C. et al. Neurobiol. Aging 57, e9–247 (2017).

25. Anderson, C.A. et al. Nat. Protoc. 5, 1564–1573 (2010).

26. Yang, J. et al. Nat. Genetics 44, 369–375 (2012).

27. Watanabe, K. et al. Nat. Commun. 8, 1826 (2017).

28. Li, H. & Durbin, R. Bioinformatics 25, 1754–1760 (2009).

29. DePristo, M.A. et al. Nat. Genetics 43, 491–498 (2011).

30. Jun, G. et al. Am. J. Hum. Genet. 91, 839–848 (2012).

31. Chang, C.C. et al. Gigascience 4, 7 (2015).

